# STARQ: Domain-Invariant Brainstem Nuclei Segmentation and Signal Quantification

**DOI:** 10.1101/2023.11.07.566040

**Authors:** Julia Kaiser, Dana Luong, Eunseo Sung, Asim Iqbal, Vibhu Sahni

**Author notes:** Corresponding authors: Asim Iqbal and Vibhu Sahni.

## Abstract

Brainstem nuclei are hard to distinguish due to very few distinctive features which makes detecting them with high accuracy extremely difficult. We introduce StARQ that builds on SeBRe, a deep learning-based framework to segment regions of interest. StARQ provides new functionalities for automated segmentation of brainstem nuclei at high granularity, and quantification of underlying neural features such as axonal tracings, and synaptic punctae. StARQ will serve as a toolbox for generalized brainstem analysis, enabling reliable high-throughput computational analysis with open-source models.

---

The evolution of deep learning applications for analyzing brain imaging data has been instrumental in advancing our understanding of brain function. These algorithms have demonstrated remarkable capabilities for analyzing brain-wide and spinal cord imaging data (Iqbal et al. [2019a], Li et al. [2023],Liang and Zhang [2023], Fiederling et al. [2021], Iqbal et al. [2019b], to mention a few), which have provided deeper insights into these neural networks. The brainstem is a key hub for processing information across multiple modalities such as motor control, sensory processing, and autonomic regulation. Current deep learning frameworks have been used to segment broad regions across the brainstem in human MRI images (Laiton-Bonadiez et al. [2022]), however their application for the identification and analysis of brainstem nuclei in 2D tissue sections has been limited. This is likely due to the absence of distinctive features that distinguish different brainstem nuclei. Therefore, there is no generalizable tool that can effectively tackle the segmentation of brainstem substructures and apply downstream analyses such as signal quantification in a high-throughput manner. To address this limitation, we provide StARQ (an extension of SeBRe - Iqbal et al. [2019a]), as a scalable machine learning framework that can accurately segment the regions of interest in the mouse brainstem and uncover signal distribution across its distinct nuclei to enable more in-depth analyses.

One of the standout features of StARQ is its domain generalization capability. It can perform region-wise segmentation with remarkable robustness. When trained on one set of source images, it can still accurately segment regions in unseen target images with scalable variational shifts caused by the scale, size, intensity. This clearly demonstrates its consistency and reliable performance even when applied to previously unseen target image domains. We open-source a model bank of highly specialized brainstem models to segment Midbrain, Pons, Medulla and their complex sub-nuclei structures as well as code in PyTorch (Paszke et al. [2017]) for the usage of pre-trained and fine-tuning of these models by the community for their customized datasets.

## StARQ framework

The block-diagram architecture of our framework is shown in **Figure 1-A**, whereas the detailed architecture of the deep neural network in shown in **Supplementary Figure 1**. In brief, to create the model that users can apply for accurate registration of brainstem regions of interest, we used coronal sections stained with a fluorescent Nissl (fNissl) that performs better in comparison with DAPI stained image sections (DICE score = 0.84 for high-level brainstem segmentation using fNissl sections and DICE score = 0.71 for DAPI stained sections). We manually annotated the training dataset of fNissl stained sections following the Reference Coronal Atlas provided by the Allen Institute (see examples in **Supplementary Figure 2, Supplementary Figure 3** and **Supplementary Figure 4**). We trained the model on brainstem sections spanning its rostro-caudal extent starting with the midbrain, through the pons and medulla. The model was trained and optimized (**Supplementary Figure 5**), and tested on a validation dataset for downstream segmentation (**Supplementary Figure 6**). This provided model now enables automated segmentation of both high-level (midbrain, pons and medulla) and low-level (brainstem nuclei within high-level regions, **Supplemental Table 1**) detection. The model also enables subsequent quantification of the signal distribution (e.g., labeled axonal collaterals) within these regions of interest in another fluorescent channel.

**Figure 1:**
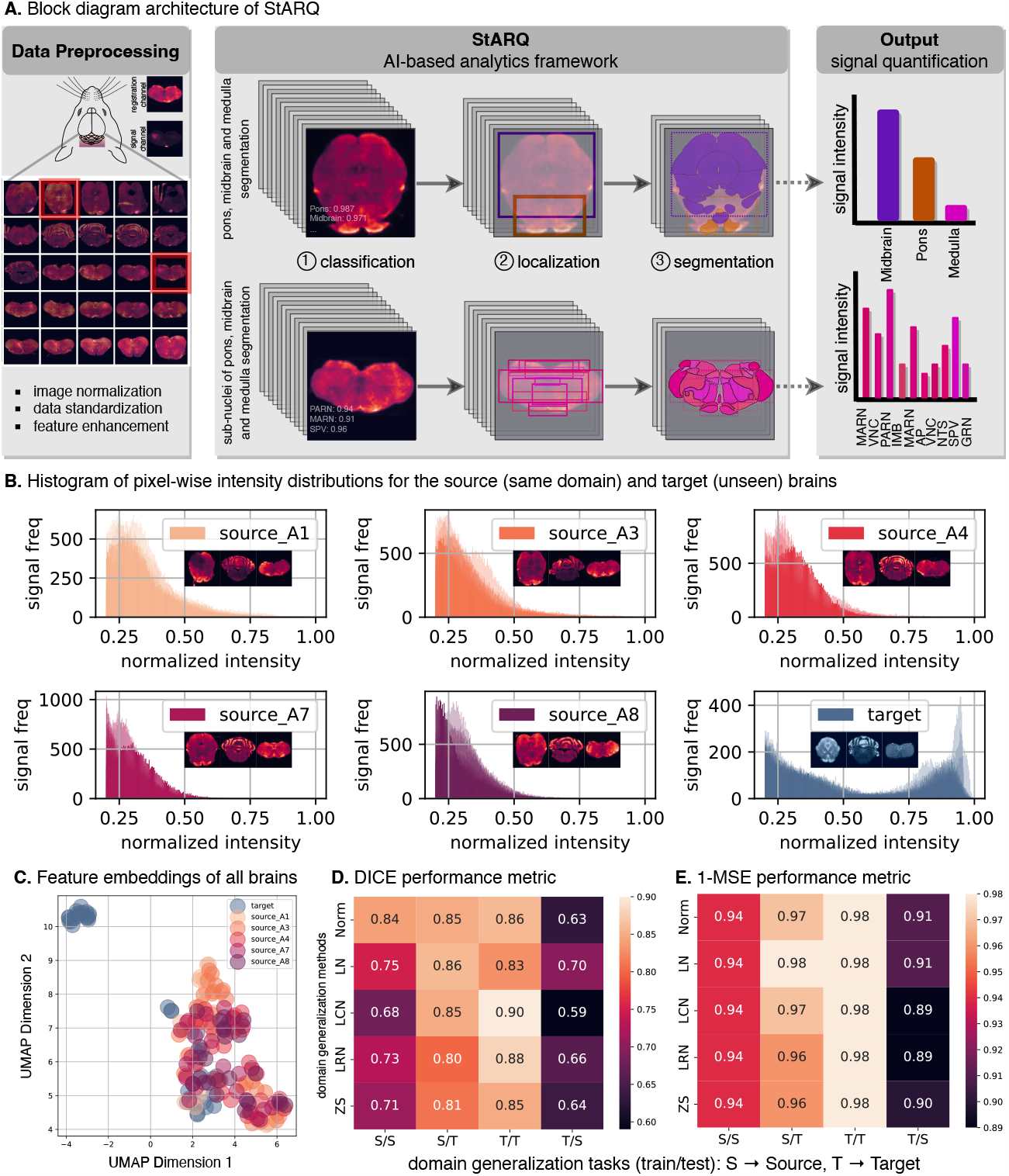
(**A**) Block diagram workflow of StARQ. The input brainstem image sections are taken through data normalization and domain-generalized feature enhancement steps. During this process, each image is also padded for maintaining image size consistency prior to feeding to the trained networks. Each image is passed through multi-stage deep neural networks, trained to segment high- (e.g. pons, midbrain, medulla) and low-level brain brainstem regions. Each network contains three heads to classify, localize and segment the regions of interest. Following this, the downstream analysis includes extraction of intensity/region in the signal channel for each brainstem image. (**B**) Domain-generalization feature of StARQ. We include powerful image feature enhancement techniques in the framework to provide robust performance for new and “unseen” datasets. Panels show the intensity-wise distribution of 5 brains with similar intensity distribution, namely “source” brains used for training, as well as one “unseen target” brain from Allen Brain database. (**C**) Panel shows the domain-shift present in the target (blue) brain as compared to source brains, visualized through low-dimensional UMAP embeddings. (**D**-**E**) shows the performance of StARQ in handling domain-shift robustness by training on ‘source-only’ and testing on ‘target’ data and vice versa. The added feature enhancement techniques in StARQ (Local Normalization, Local Contrast Normalization, Local Response Normalization and Z-Score) are shown improving the performance on unseen target set for brainstem segmentation.

## Domain generalized feature enhancement in StARQ

For most experimental data that requires image acquisition, significant variation arises from differences in imaging set ups (e.g. different microscopes), staining protocols, tissue folding, etc. To tackle these issues of shifts in image domains arising from such variables, we introduce a feature of domain generalization in StARQ. This is enabled by training the models on feature representations, captured by introducing powerful image preprocessing techniques such as Local Normalization (LN - Sage and Unser [2003]), Local Contrast Normalization (LCN - Jarrett et al. [2009]), Local Response Normalization (LRN - Krizhevsky et al. [2012]) and Z-Score (Huck et al. [1986]). These pixel normalization techniques enhance the high-frequency pixel-based features in the input image while suppressing the background noise (**Supplementary Figure 7**)

To handle the domain shift robustness in the brainstem imaging datasets, we first computed the intensity-based distribution (histogram) of five source brains, as shown in **Figure 1-B**. The intensity distribution plot for all pixels across all these brains shows that they all follow a similar trend i.e. skewed right. We computed the same distribution in a sample target brainstem image, from the Allen Brain Atlas, which was acquired using a different acquisition set up. The intensity distribution in this target brain follows a bimodal distribution with domain shift into the high-intensity range. Training the model on the source brain and testing on such target data with a very different intensity distribution poses a challenge to the model. Since the training data is not drawn from the same distribution with pre-existing inherent variational shifts as compared to the unseen testing (target) data and vice versa, this further challenges the model. This is also visualized by UMAP feature embeddings of all the brains in **Figure 1-C**. To overcome this challenge we introduced domain shift robustness for the downstream task of segmenting high-level brainstem regions. To enhance the segmentation performance of StARQ on domain generalization tasks, we elect the aforementioned domain normalization techniques (LN, LCN, LRN and Z-Score). Qualitative examples of domain adaptation by the above-listed techniques, across different brain markers, are shown in **Supplementary Figure 7**. It can be observed that these techniques tend to minimize the distributional shift across the different imaging domains and highlight the generalized features of brain regions (including cortex). LN is observed to filter out the low frequency pixels whereas LRN focuses on enhancing contrastive features.

We evaluated the effects of adding domain generalized feature processing in the model pre-processing stage. These adaptive techniques are typically applied during image processing, in order to rectify a distribution shift between the source and target domains, and to increase domain homogeneity in the data. This step allows downstream image-analysis and image-manipulation tasks to be performed on the pre-treated dataset in a domain invariant manner. We ran a series of experiments by training the model on a source (S) brain and observed its segmentation performance on the target (T) brain by applying the pre-processing from different image normalization techniques. The results in **Figure 1-D** shows the DICE scores of each combination of experiments. As expected, the results for the S/S show high-performance for the raw data distribution. Encouragingly, the S/T domain shift shows high-performing results for both LN and LCN as well. In contrast, T/T showed high-performing segmentation after applying LCN followed by LRN. We also performed the converse (T/S), where we trained the model on the T brain and tested on S. For this, application of LN and LRN provided the best segmentation results. Overall a similar trend is observed on the 1-MSE metric in **Figure 1-E** with less sensitivity as compared to the DICE metric.

These results show that the selection of domain generalization features in StARQ can have a drastic impact on the segmentation of brainstem regions, especially in instances where the training and validation datasets are drawn from different distributions. We provide the end users access to these multiple preprocessing features in StARQ to test for optimal use and applicability in their custom datasets.

## StARQ-based brainstem data analytics

To demonstrate the effectiveness of utilizing StARQ for performing high-throughput brainstem imaging analysis, we used coronal brainstem sections from mice in which we had injected an AAV-eGFP reporter into the cerebral cortex to visualize cortical axonal projections to the brainstem. While motor cortical input into the brainstem has been well studied (Kuypers [1982], Armand [1982], CANEDO [1997]), the brainstem innervation from other sensorimotor regions is less clear. We thus injected the AAV-GFP reporter across a large area of the lateral sensorimotor cortex to label a broad population of subcerebral projections and identify brainstem regions that receive inputs from this cortical region and regions that do not.

We used StARQ to perform segmentation, using fNissl (segmentation channel) of the high- and low-level brainstem regions and then applied signal quantification (signal channel) to investigate the distribution of axonal collaterals across the distinct brainstem nuclei. Our results in **Figure 2-A** and **Figure 2-B** demonstrate that StARQ can perform accurate registration of distinct brainstem nuclei in midbrain, pons, and medulla, with high confidence. The heatmap (**Figure 2-B**) shows that the highest distribution of the signal (i.e. axonal collaterals within each nucleus) are consistent across 3 mice. Our investigation demonstrates a strong connection between the sensorimotor cortex to all three high-level regions of this study (midbrain, pons, medulla) (Peng et al. [2021]), consistent with previous work using sparse labeling and whole-brain imaging. Within the midbrain, and consistent with previous reports, we find stronger connections to SNr than TRN or PAG. Within the pons, the strongest connections are with the PG, with less connections to PRN and TRN and almost no connections to PB. Inputs into medulla seem to vary the most across cell-types and regions. In our study, with most of the lateral sensorimotor cortex labeled, we corroborate previous findings of stronger sensorimotor cortical input into PARN, MDRNd, SPC, as well as lesser input into Cu and Ecu (Yang et al. [2023], Peng et al. [2021]). The LDT (lateral dorsal tegmental nucleus) and LC (locus coeruleus) in the pons, and AP (area postrema) in the medulla do not receive inputs from the lateral sensorimotor cortex. This provides an automated map of connectivity between the lateral sensorimotor cortex and the brainstem.

**Figure 2:**
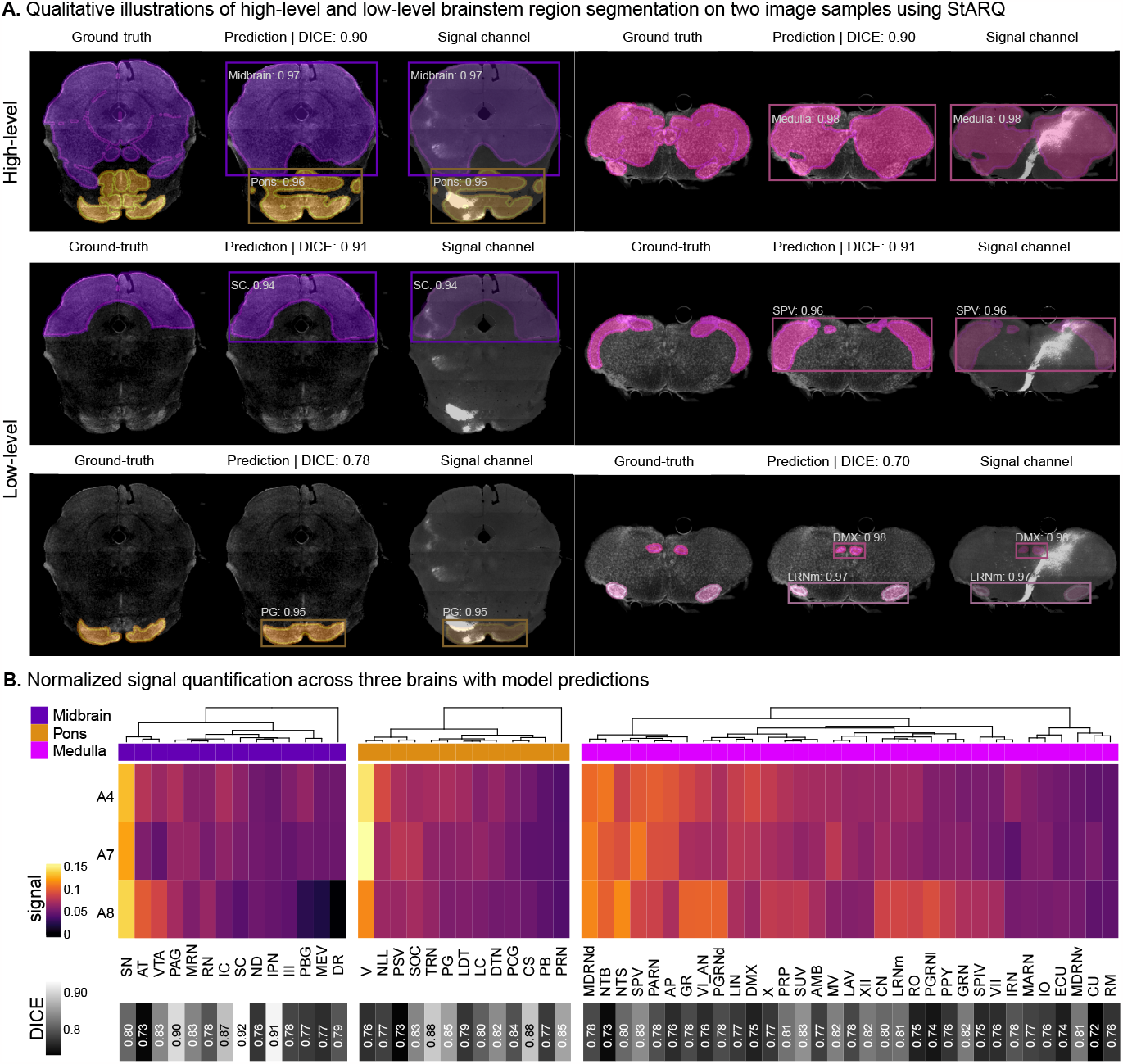
Qualitative and quantitative performance of StARQ on “unseen” validation dataset. **A** demonstrates the ground-truth and model predictions (overlaid on fNissl and signal channel) on two brainstem images for high-level (midbrain, pons and medulla) and a few low-level segmentation of nuclie. **B** shows the signal intensity measurement across three (A4,A7,A8) validation brains with model prediction alignment score (DICE) with ground-truth for each segmentated nuclie in the bottom.

In this study, we assessed axonal projections from a large area of the sensorimotor cortex to demonstrate effectiveness of StARQ for these kinds of analyses. However, previous reports highlight the importance of region and cell-type specificity to function, even within a relatively clearly organized structure such as the cortex (Tasic et al. [2018], Saunders et al. [2018], Economo et al. [2018]). In future studies, StARQ can be utilized to systematically assess brainstem innervation from different brain regions, including but not limited to the cortex, using reporter mouse lines, and different injection sites in both healthy mice and in mouse models of neurological disorders.

## StARQ brainstem region-specific model bank

In StARQ, we offer a range of specialized models (**Supplemental Table 1**). The first high-level model segments parent regions such as Midbrain, Pons and Medulla. Within these three parent categories, the Midbrain_Sensory model is designed to segment SC, IC, PBG, and MEV regions, making it tailored for exploring sensory processing in the midbrain. For motor control investigations, the Midbrain_Motor model is trained to segment PAG, III, MRN, SN, RN, and AT regions. In behavioral region analytics, the Midbrain_Behavioral model is tailored for segmenting IPN, DR, VTA, and ND regions. In the Pons, similar to the Midbrain, we provide sensory and motor region models. The Pons_Sensory model precisely segments PB, SOC, NLL, and PSV regions, while the Pons_Motor model is designed for segmenting PG, TRN, V, PCG, and DTN regions. Pons_Behavioral model caters to CS, PRN, LDT, and LC regions. Lastly, the Medulla_Sensory model is tailored to segment CN, SPV, CU, NTS, ECU, AP, and GR regions. The Medulla_Motor models encompass XII, VII, MARN, AMB, and IO regions, while the Medulla_Behavioral category includes models for segmenting RM and RO regions. End users can select the high-level (Midbrain, Pons, Medulla) as well as low-level region specialized model of interest from the drop-down menu selection in the framework (**Supplemental Figure 8**).

In summary, StARQ is a truly versatile and accurate solution for mouse brainstem image analysis. Our results show that it overcomes the necessary challenges in automated segmentation of brainstem regions. This tool will have broad applicability across multiple anatomical investigations of the brainstem. StARQ will not only help advance our understanding of normal function but also aid in the investigation of neurological disorders, where precise segmentation of brainstem structures could identify novel circuit substrates of dysfunction and also improve monitoring of disease progression.

## Methods

### StARQ deep learning model architecture

StARQ was designed by optimization of Mask R-CNN architecture, and constructed by using a convolutional backbone that comprises the first five stages of the very deep ResNet-101 and FPN architectures (our network architecture is shown in **Supplementary Figure 1**). The feature map is processed by an RPN, which applies a convolutional neural network over the feature map in a sliding-window fashion. The RPN segregates and forwards the predicted n potential regions of interest (ROI) from each window to the Mask R-CNN (He et al. [2017]) ‘heads’ on the basis of the FPN. The ROI feature maps undergo a critical feature pooling operation by a pyramidal ROIAlign layer, which preserves a pixel-wise correspondence to the original image. Each level of the pyramidal ROIAlign layer is assigned an ROI feature map from the different levels of the FPN backbone (depending on the feature map area), which returns n pooled feature maps. Three arms of the FPN perform the core operations of brain region segmentation. The ‘classifier’ and ‘regressor’ heads—inherited from the Faster R-CNN (Girshick [2015]) and identify distinct brain regions and compute region-specific bounding boxes. The classifier output layer returns a discrete probability distribution for different object classes. The regressor output layer gives the four (x coordinate, y coordinate, width, height) bounding-box regression offsets to be applied for each class. A fully convolutional network forms the more recent mask-prediction arm, which returns a binary mask spanning each segmented brain region. The network is trained using a stochastic gradient descent algorithm that minimizes a multi-task loss (L) corresponding to each labelled ROI. Following describes the individual losses and **Supplementary Figure 5** shows these loss curves for a sample high-level and low-level model training:

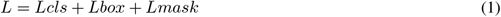

where Lcls, Lreg and Lmask are the region classification, bounding box regression and predicted masks’ loss, respectively, as defined below:

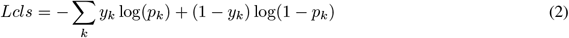

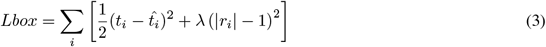

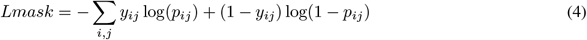

where *p*_*k*_ is the probability of the kth proposed ROI (Region of Interest or anchor) and *y*_*k*_ is the ground-truth class labels. The loss is computed for each class (k) and summer over all losses. We use the cross-entropy loss, which penalizes the model more when it predicts the wrong class with high confidence. The vector *t*_*i*_ represents the four coordinates that characterize the predicted anchor bounding box, whereas vector 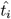 represents the coordinates for the ground-truth box that correspond to a positive anchor. The loss has two components: the first component computes the squared difference between predicted and ground truth coordinates, normalized by a factor of 1/2. The second component penalizes the deviation of the predicted box size (|*r*_*i*_|) from a predefined “anchor” box size. This term encourages the model to predict bounding boxes with a consistent aspect ratio and the loss is summed over all instances (i) in the dataset.

The Lcls for each anchor is calculated as logarithmic loss for class labels. The Lreg term is a regression loss function robust and parameters ncls and nreg are the normalizations for classification and regression losses, respectively, weighted by a balancing parameter, *µ*. The Lmask term is computed as average cross-entropy loss for per-pixel binary classification (mask probabilities (*p*_*ij*_) and ground truth binary masks (*y*_*ij*_)), applied to each ROI.

### StARQ configuration

The implementation of StARQ generally follows the original work in SeBRe (Iqbal et al. [2019a]), with limited hyperparameter optimization for the brainstem section dataset. Training on the brain section dataset is initialized with pre-trained weights. Each batch slice consists of a single brainstem section image per graphics-processing unit; batch normalization layers are inactivated to optimize training for the small effective batch size. Training is performed using an NVIDIA Tesla T4 GPU (graphics-processing unit). The training regime comprises two stages. First, the network heads are trained for 5,000 iterations at a learning rate of 0.001 and learning momentum of 0.9. Second, all of the layers are fine tuned for 5,000 iterations at a reduced learning rate of 0.0001. During inference, diverging from the original model, the mask branch is applied to the highest scoring detection boxes (limited to the unique number of region a model can detect), proposed by the RPN. The maximum number of ground-truth instances detected per image is also limited to the unique number of regions that the model can segment (to avoid erroneous duplicate instances of region-specific masks). Adopting a more stringent approach, the minimum probability threshold for instance detection is raised to 0.9 to improve the accuracy of instance segmentation, while this can be adjusted based on user’s choice.

### Mice

Wild-type CD1 mice were obtained from Charles River Laboratories (Wilmington, MA). The day of birth was designated at P0 (postnatal day 0). All mouse studies were performed in accordance with institutional and federal guidelines and were approved by the Weill Cornell Medical College Institutional animal care and use committee.

### Experimental data collection

Subcerebral projection neurons (SCPN) were labeled at P1 by injection of AAV1-hSyn-eGFP (2.7 × 10^13^ GC/mL) using ultrasound-guided backscatter microscopy (Vevo 2100; VisualSonics, Toronto, Canada) via a pulled glass micropipette with a nanojector (Nanoject II, Drummond Scientific, Broomall, PA) as previously described in Sahni et al. [2021a] and Sahni et al. [2021b]. At P28, brains were dissected, post-fixed in 4% PFA overnight, and stored at 4°C in PBS until further use. Brainstem tissue was cryopreserved in 30% sucrose overnight and frozen in Tissue-Tek OCT Compound (Sakura Finetek, Torrance, CA). Serial coronal brainstem sections (50 *µ*m) were prepared on a cryostat, and every 3rd tissue section was selected for use in the StARQ pipeline. Sections were labeled free-floating with NeuroTrace™ 640/660 Deep-Red Fluorescent Nissl Stain (Invitrogen, Carlsbad, CA) and mounted and coverslipped using DAPI-Fluoromount-G (VWR). Slices were imaged at 10x on a Zeiss Axioimager M2 microscope using the Stereo Investigator software (MBF Biosciences).

### Brainstem imaging dataset and preprocessing

To ensure data diversity and robust model training, we employed a total of five brains with total 120 sections covering 52 unique regions (nuclei of brain-stem parent regions i.e. midbrain, pons and medulla) meticulously annotated by three expert human annotators. Out of these, two brains were designated for training and augmentation (see examples in **Supplementary Figure 3** and **Supplementary Figure 4**), while the remaining three were reserved for testing (see examples in **Supplementary Figure 9** and **Supplementary Figure 10**.

For training data, we introduced augmentation by applying rotations within the range of -10 to 10 degrees to each section with a sampling size of 2 degrees (see examples in **Supplementary Figure 11**), enhancing the dataset’s variability, resulting in 540 images with ground-truth labels as total size of the training data. This augmentation is crucial for improving the model’s ability to handle diverse orientations of brainstem sections. Prior to model training and testing, all brainstem section images were preprocessed to ensure uniformity and normalization. Specifically, we mapped all images to a consistent size and standardized the format, and then padded them with an empty background. This padding is essential to maintain uniform dimensions and ensure compatibility during the training and testing of the deep neural network.

The pre-processing step can be represented mathematically as follows, where *I*_original_ denotes the input section before padding, *I*_padded_ represents the section after padding, *w* and *h* are the desired width and height of the padded section, and Pad represents the padding operation:

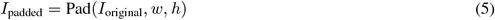

### Allen brain imaging dataset

The Allen Brain Reference atlas (atlas.brain-map.org) comprises a total of 38 brain sections obtained from a single brain, processed through fNissl staining. These sections collectively cover 52 unique regions, representing nuclei of the brainstem’s parent regions, including midbrain, pons, and medulla, as annotated by expert annotators at Allen Brain Institute.

### DICE score as a quantitative metric

The DICE score is a widely-used quantitative metric in the field of image segmentation and medical imaging analysis. It provides a measure of the spatial overlap between the predicted and ground-truth segmentation masks. The DICE score is particularly useful for evaluating the accuracy and quality of segmentation results, as it takes into account both true positives (correctly segmented pixels) and the sizes of the segmented regions. The DICE score is defined by the following equation:

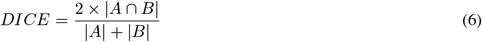

Here, *A* represents the segmented region obtained from the model’s predictions and *B* represents the ground-truth segmented region. The DICE score ranges from 0 to 1, where a score of 0 indicates no spatial overlap between the predicted and ground-truth regions, and a score of 1 implies a perfect match. In practice, DICE scores closer to 1 indicate a higher level of accuracy and similarity between the segmentation results and the true regions of interest.

### 1-MSE score as a quantitative metric

The 1-MSE (1-Mean Squared Error) score is a valuable quantitative metric employed to assess the accuracy and quality of various predictive models. It measures the dissimilarity between predicted values and true ground-truth values, with a focus on minimizing errors. The 1-MSE score is a useful tool for evaluating image similarity, here in our case it is used to measure the similarity between ground-truth and predicted masks:

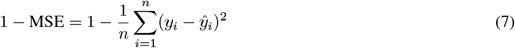

Here, *n* represents the total number of data points, (*y*_*i*_) denotes the true or ground-truth value for data point *i* and (*ŷ*_*i*_) represents the predicted value for data point *i*. The 1-MSE score ranges from 0 to 1, where a score of 1 signifies a perfect prediction with no errors, while a score of 0 suggests that the model’s predictions are as poor as simply taking the mean of the true values. In practice, a higher 1-MSE score indicates a more accurate and reliable predictive model, while a lower score implies a less precise fit.

### Signal quantification

Utilising the model bank provided with StARQ framework can automatically segment the brainstem regions of interest. After segmentation from fNissl channel, the segmented mask is applied to fetch the signal distribution in the signal channel under the masked area. The computed signal is normalised by the area of the region to measure the normalised signal distribution. Given *R*_*i*_ is an image mask for a brainstem nuclei, each brain region (*R*_*i*_) has a unique RGB color code, brain regions (RN) are filtered by their respective RGB codes and temporarily stored in the form of binary images where active *R*_*i*_ pixels are 1s, and 0s otherwise:

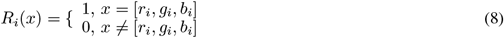

Afterwards, signal density (*S*_*i*_) of a particular brain region (*R*_*i*_) can be calculated by computing its dot product with the signal image of a given brainstem section (*A*_*i*_) and normalizing it by dividing with the area of the brain region, |*R*_*i*_|.

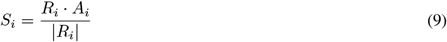

## Code availability

The source code of running a pre-trained model from the brainstem model bank as well as finetuning it on custom dataset is provided on GitHub (https://github.com/itsasimiqbal/StARQ).

---

**Supplementary Figure 1:**
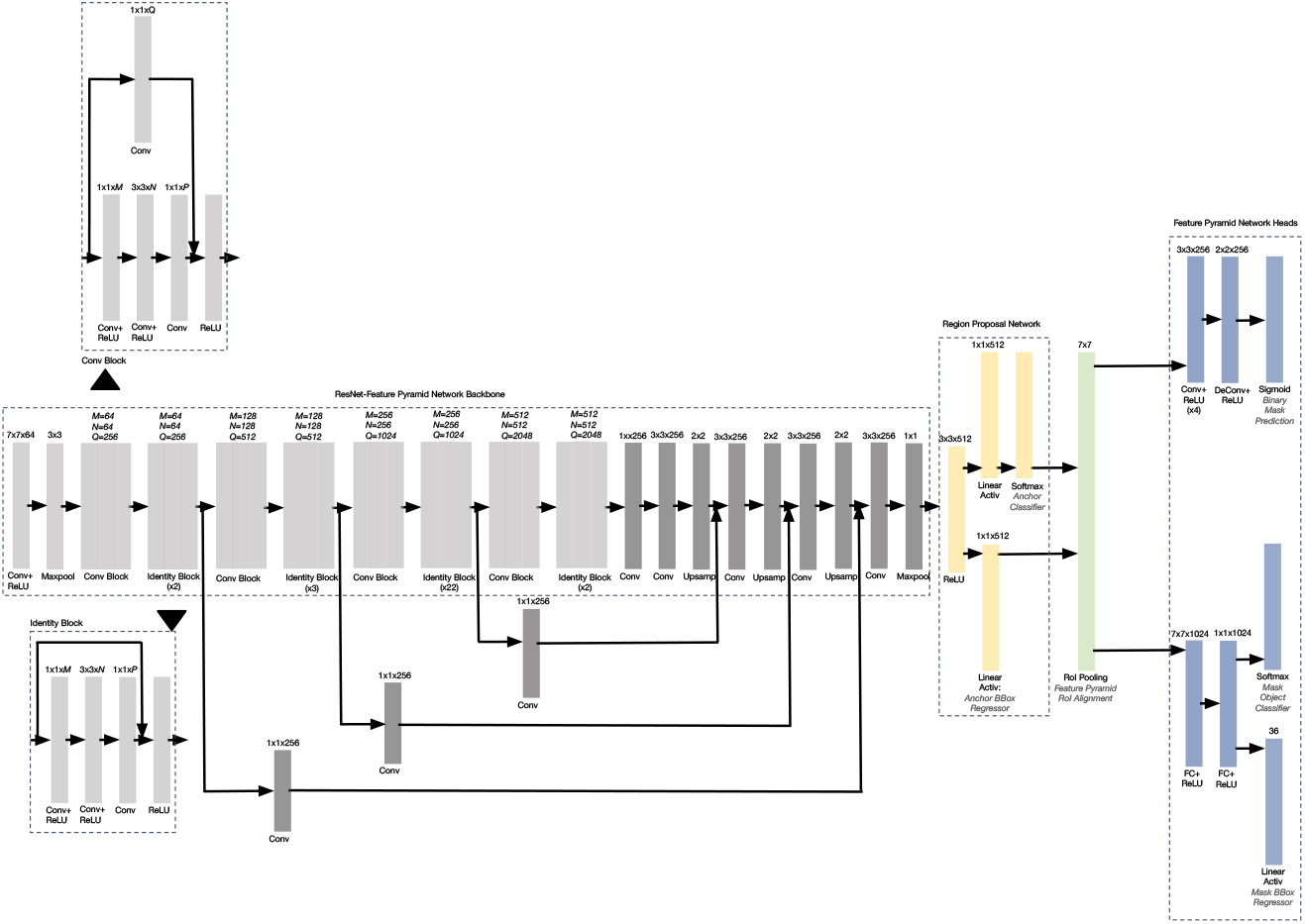
Architecture of the Mask-RNN for instance segmentation. The backbone consists of repeating convolutional blocks and identity blocks based on a ResNet-Feature Pyramid Network. The network head contains components for region proposal, feature pyramid construction, ROI pooling/alignment, and mask/bounding box prediction. Convolutional blocks downsample features using strided convolutions while identity blocks preserve feature resolution. The feature pyramid and upsampling components enable segmentation at multiple scales. ROI alignment enables precise segmentation even for small regions. The network is trained end-to-end for simultaneous detection and segmentation.

**Supplementary Figure 2:**
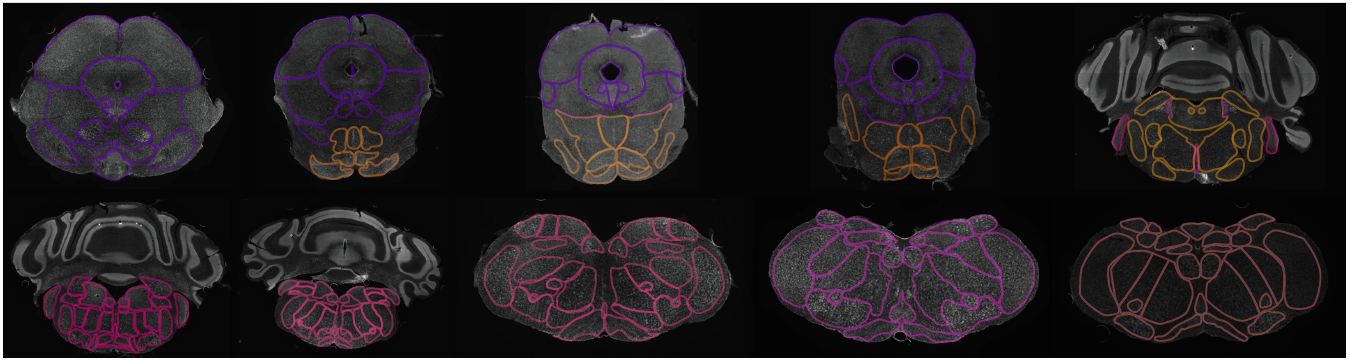
Illustration of 10 example sections covering midbrain, pons and medulla with their sub-nuclei as finely annotated by human experts.

**Supplementary Table 1:**
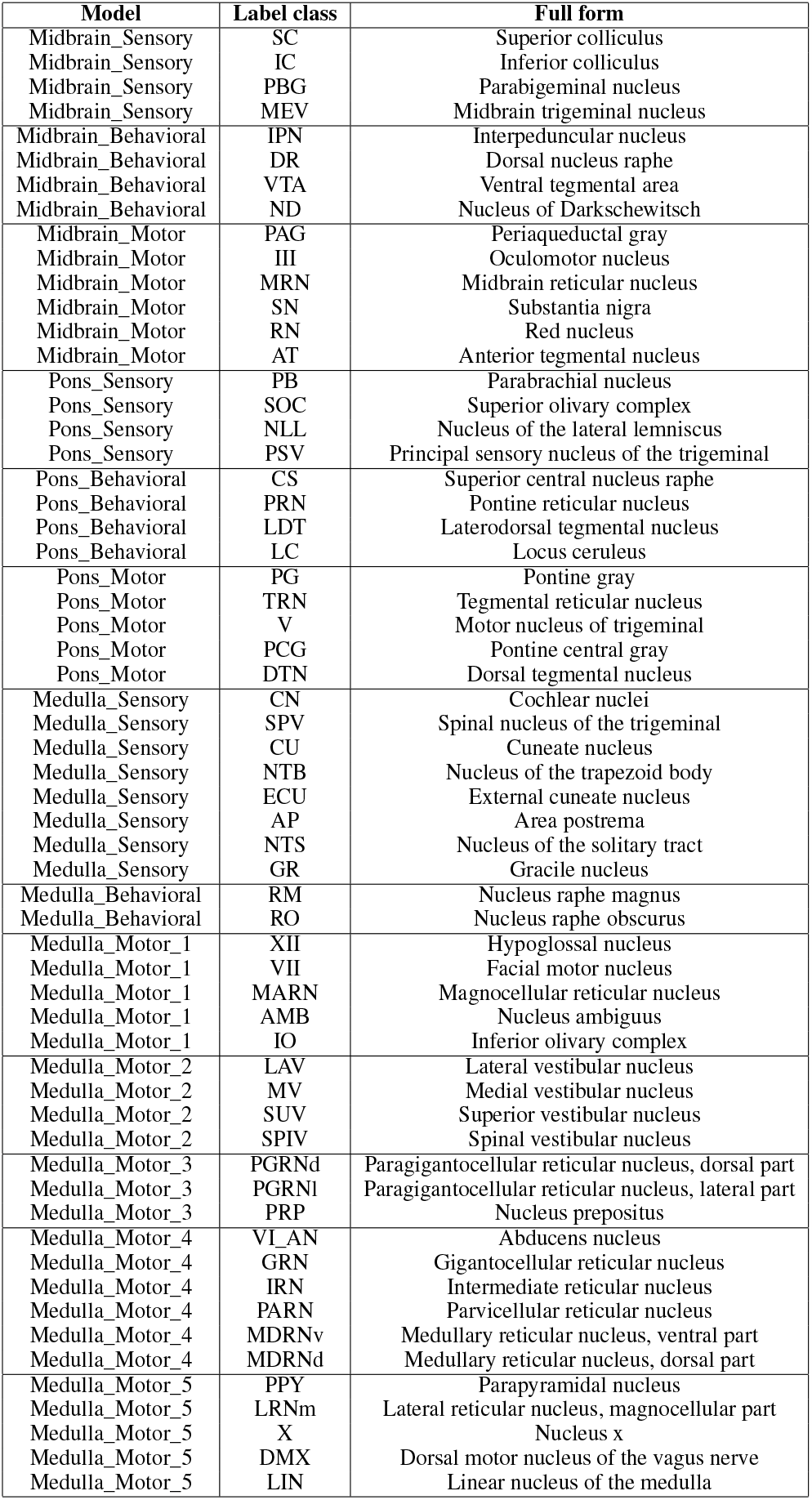
Pre-trained models and labelled classes for brainstem sub-regions.

**Supplementary Figure 3:**
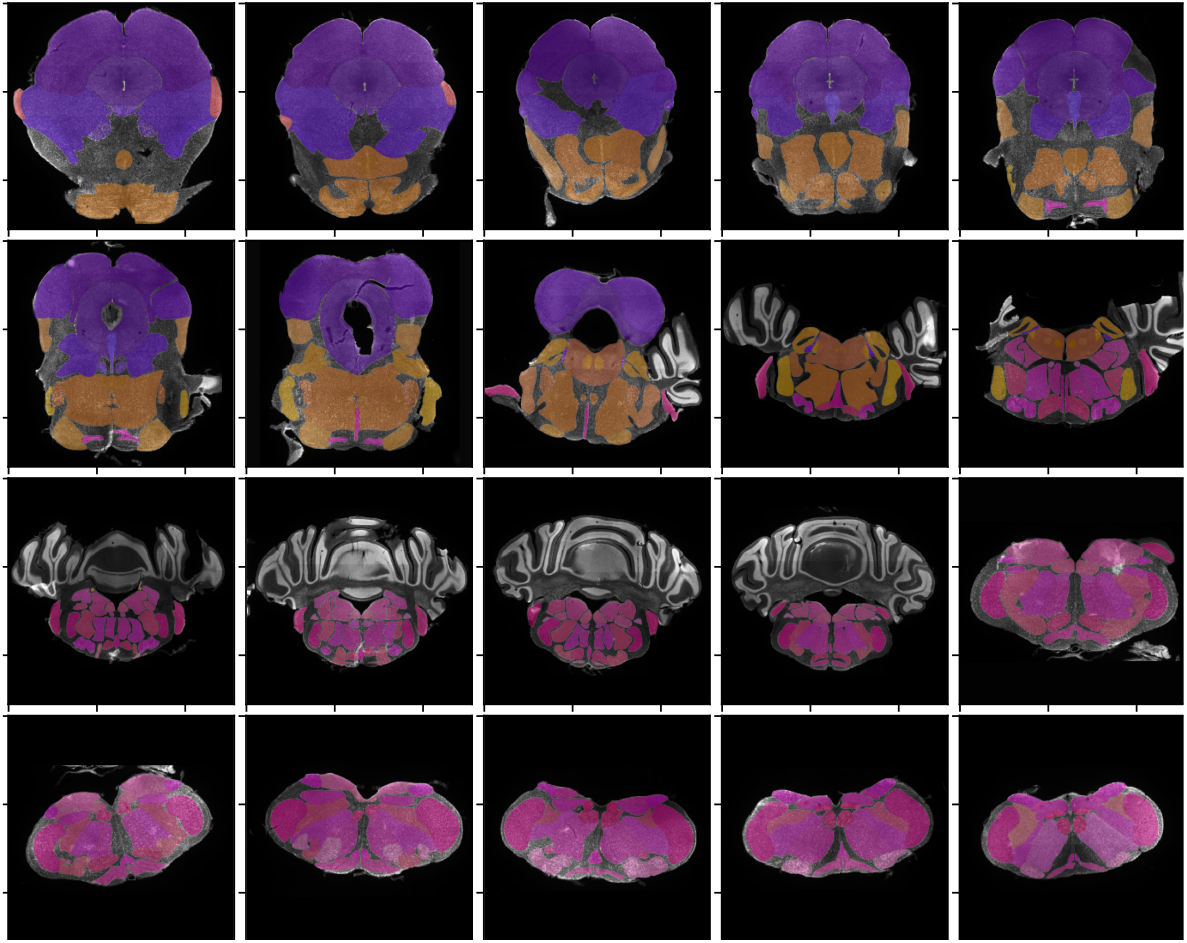
Sample of ground-annotated masks overlaid on fNissl image section drawn from a sample brain A1 used for training. Brainstem sections spanning midbrain, pons and medulla with their sub-nuclei regions are assigned a unique color code.

**Supplementary Figure 4:**
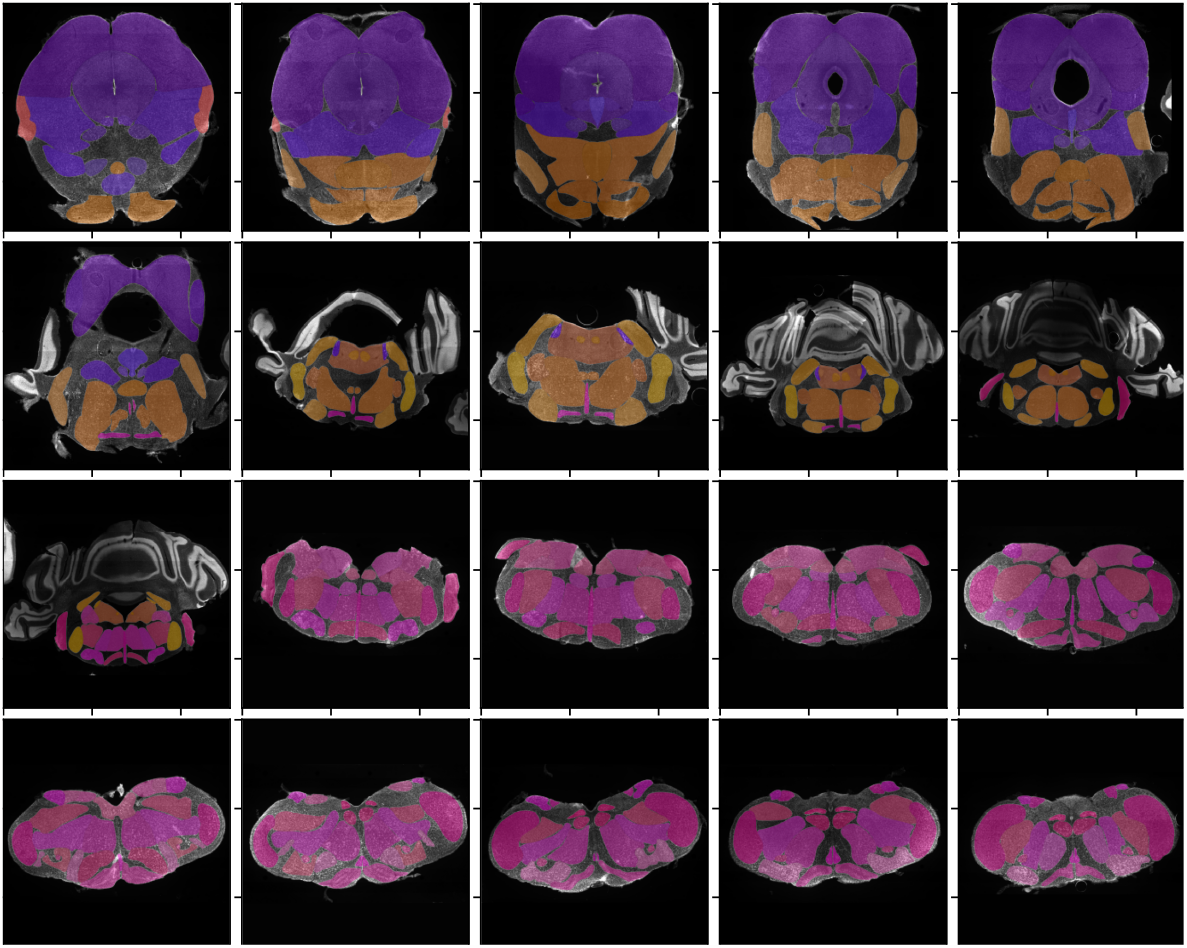
Sample of ground-annotated masks overlaid on fNissl image section drawn from a sample brain A3 used for training. Brainstem sections spanning midbrain, pons and medulla with their sub-nuclei regions are assigned a unique color code.

**Supplementary Figure 5:**
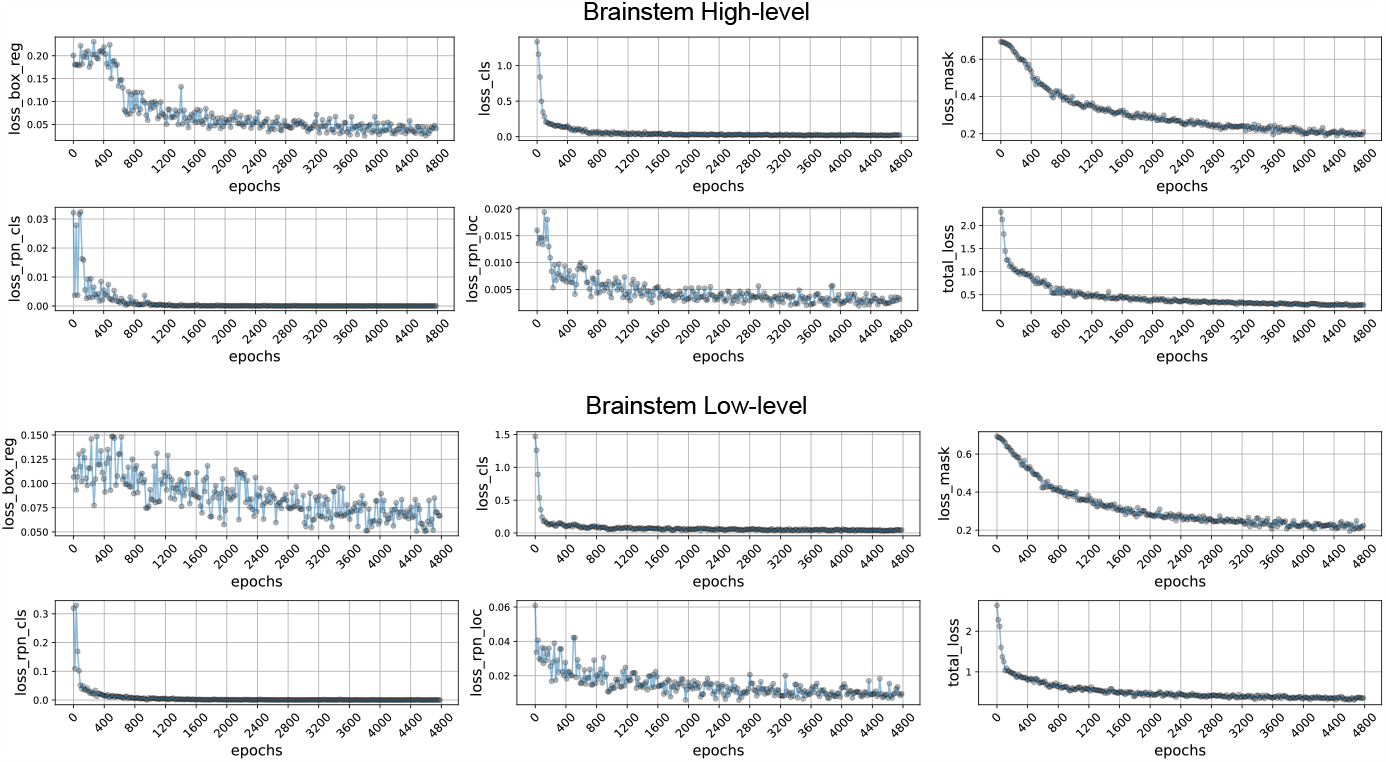
The loss curves of the high-level and one of the low-level segmentation models are shown. The losses demonstrate the classification, regression, segmentation and total combined loss. Models are trained and optimized on 5000 epochs.

**Supplementary Figure 6:**
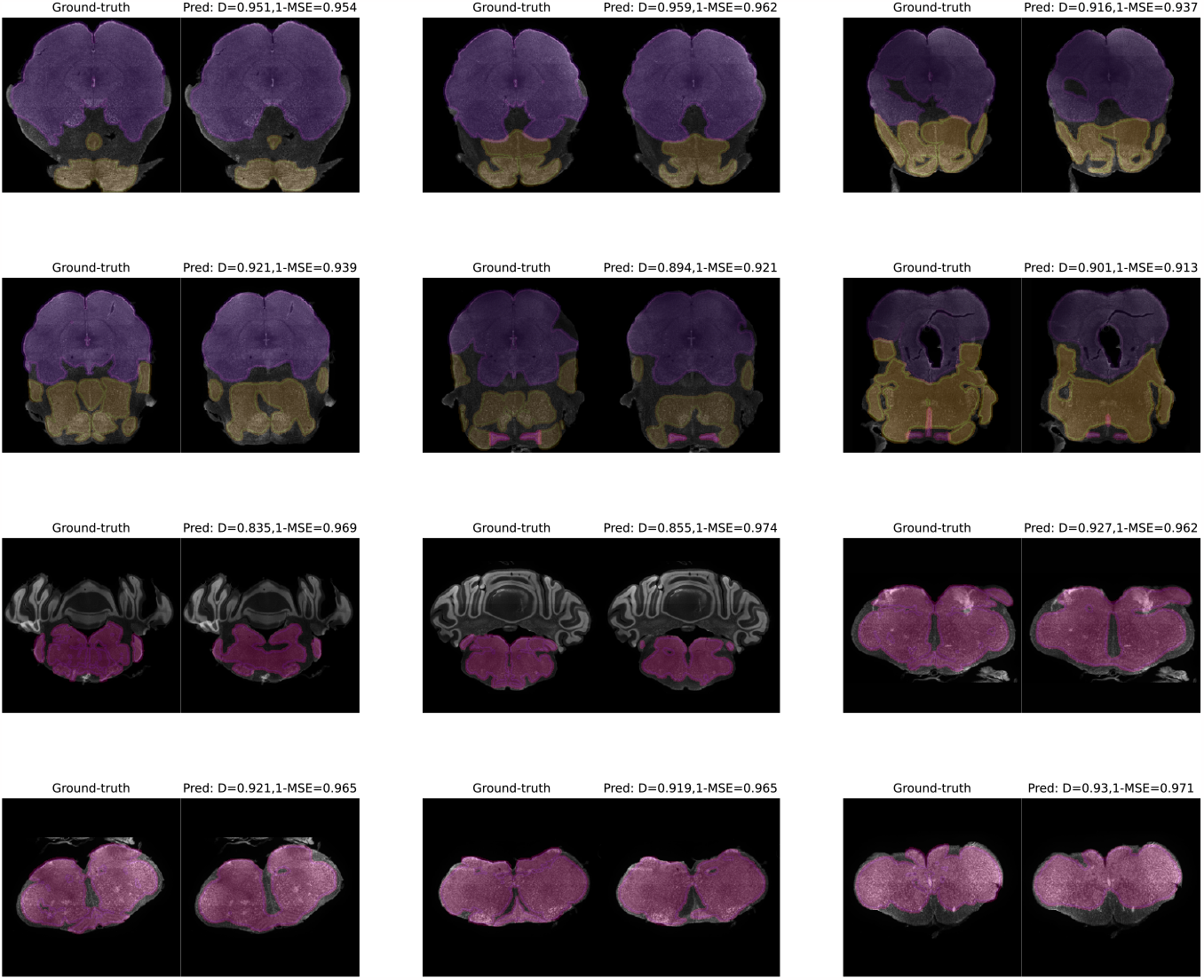
Qualitative results of the trained high-level (midbrain, pons and medulla) model on a few brainstem validation sections are shown, overlaid on the fNissl sections on which the model is trained.

**Supplementary Figure 7:**
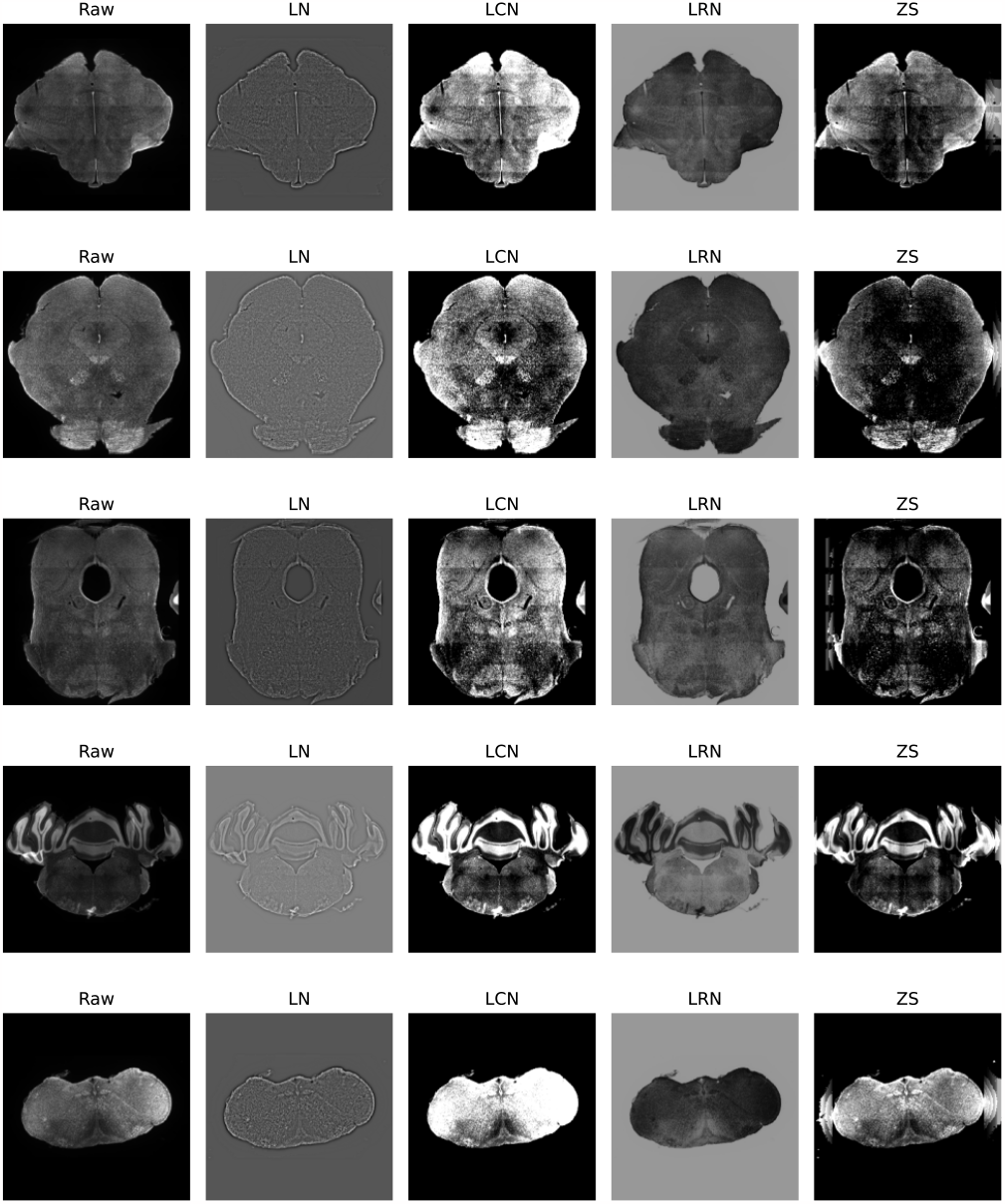
Sample qualitative illustrations by passing the original (raw) brainstem sections to domain generalization image preprocessing techniques: Local Normalization (LN - Sage and Unser [2003]), Local Contrast Normalization (LCN - Jarrett et al. [2009]), Local Response Normalization (LRN - Krizhevsky et al. [2012]) and Z-Score (Huck et al. [1986]).

**Supplementary Figure 8:**
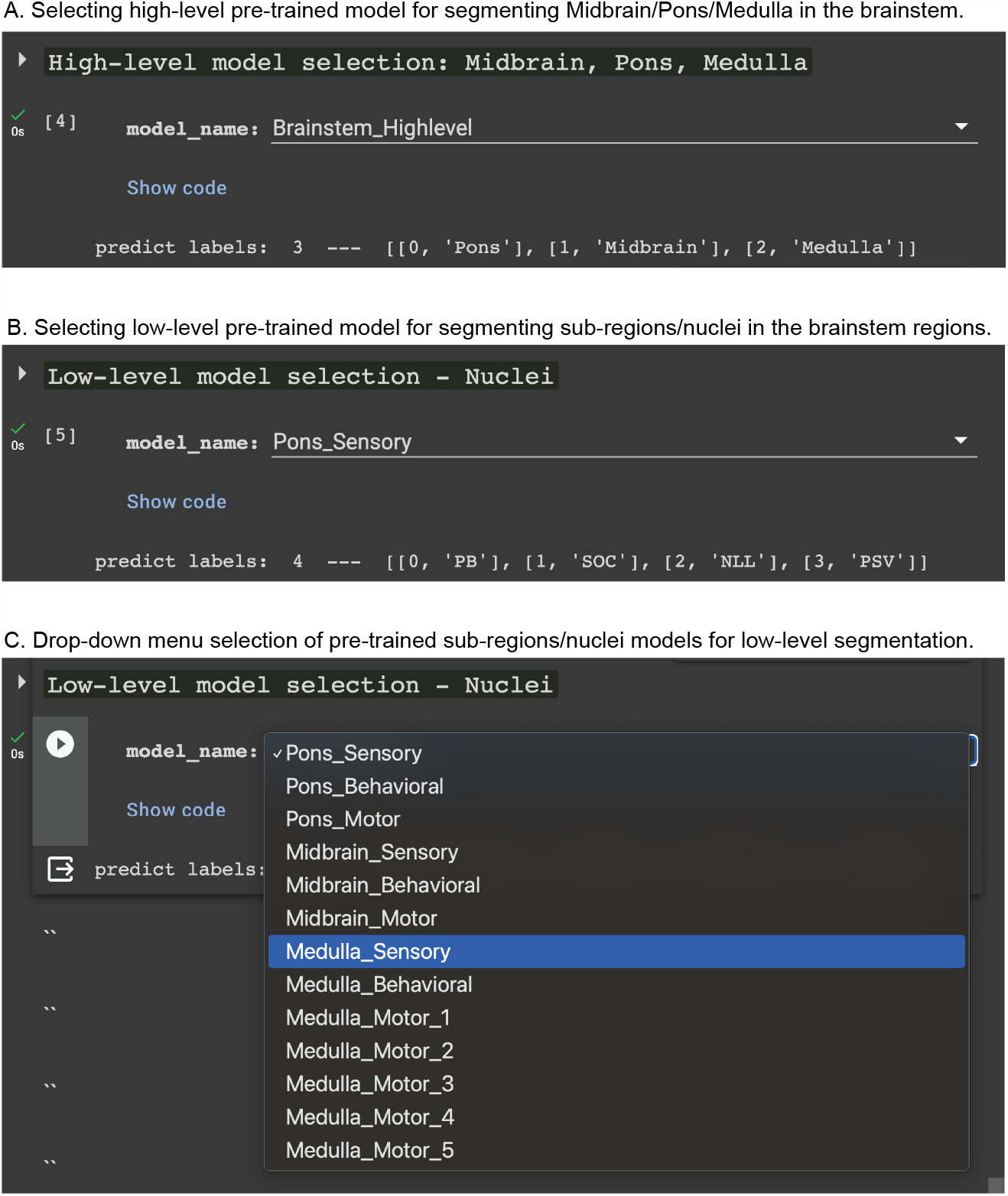
Model selection (high-level: midbrain, pons and medulla) and specialised low-level nuclei segmentation) from a drop down menu provided to faciliate the user in running their models of interest on their dataset.

**Supplementary Figure 9:**
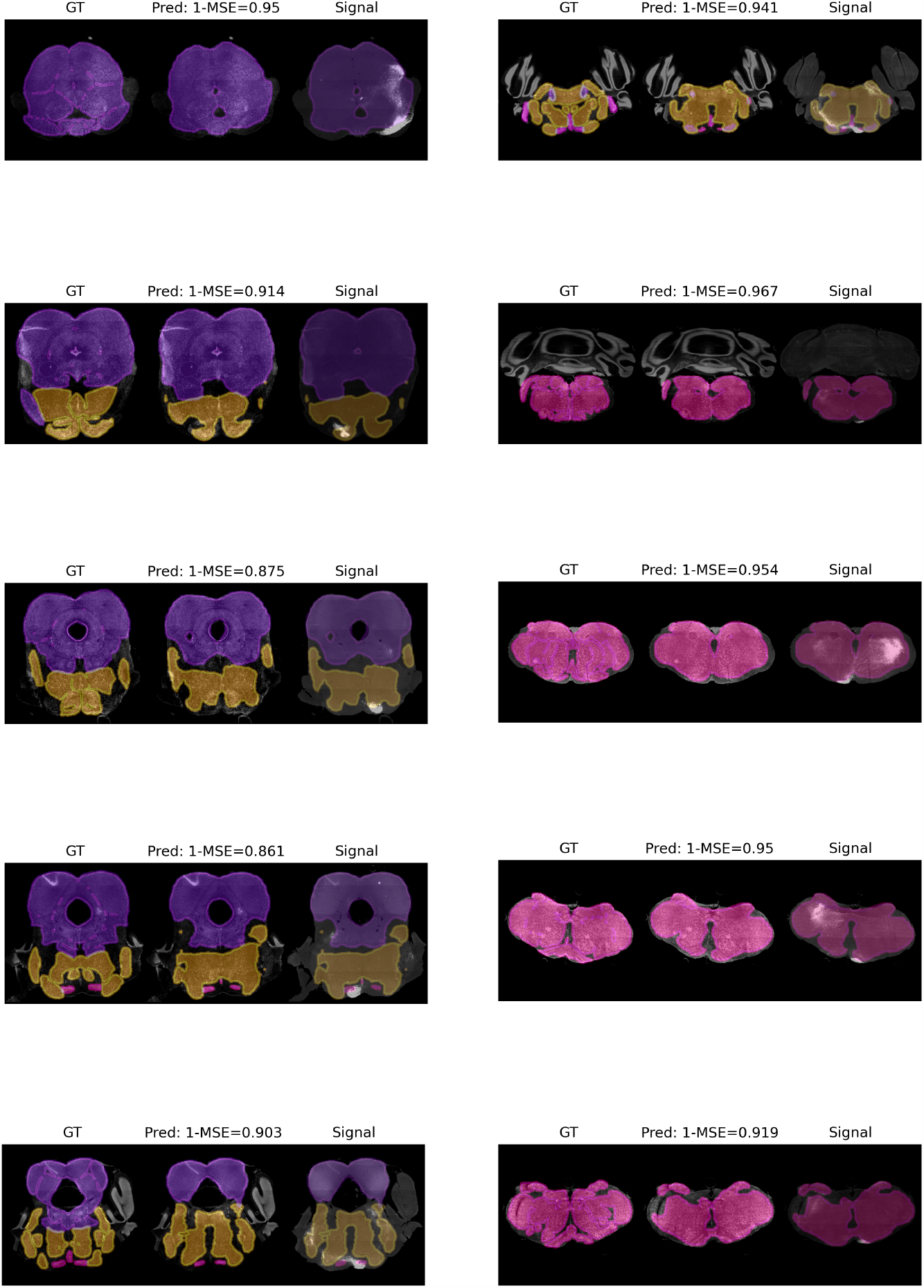
Qualitative samples of high-level (Midbrains, Pons and Medulla) brainstem region segmentation with signal overlay (GT: Ground-Truth).

**Supplementary Figure 10:**
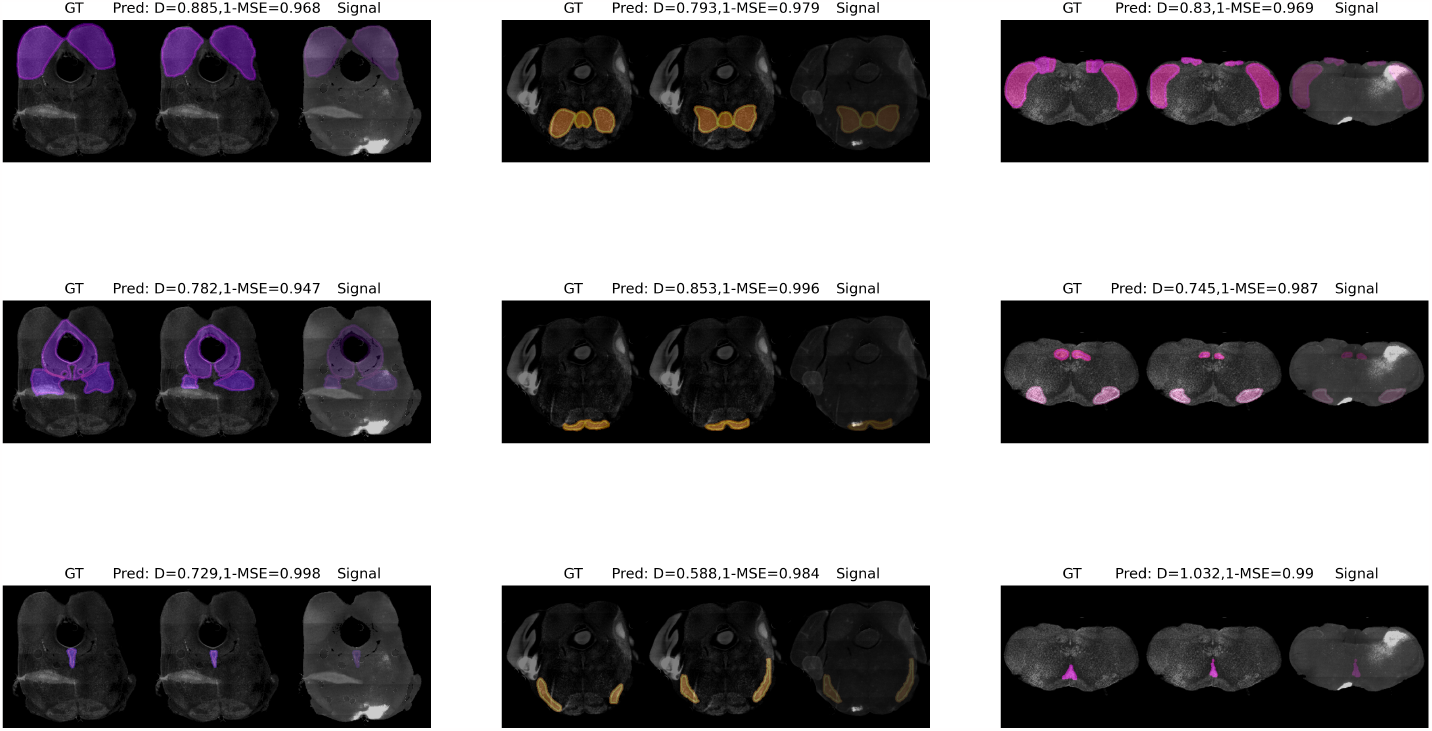
Qualitative samples of high-level (Midbrains, Pons and Medulla) brainstem region segmentation with signal overlay (GT: Ground-Truth).

**Supplementary Figure 11:**
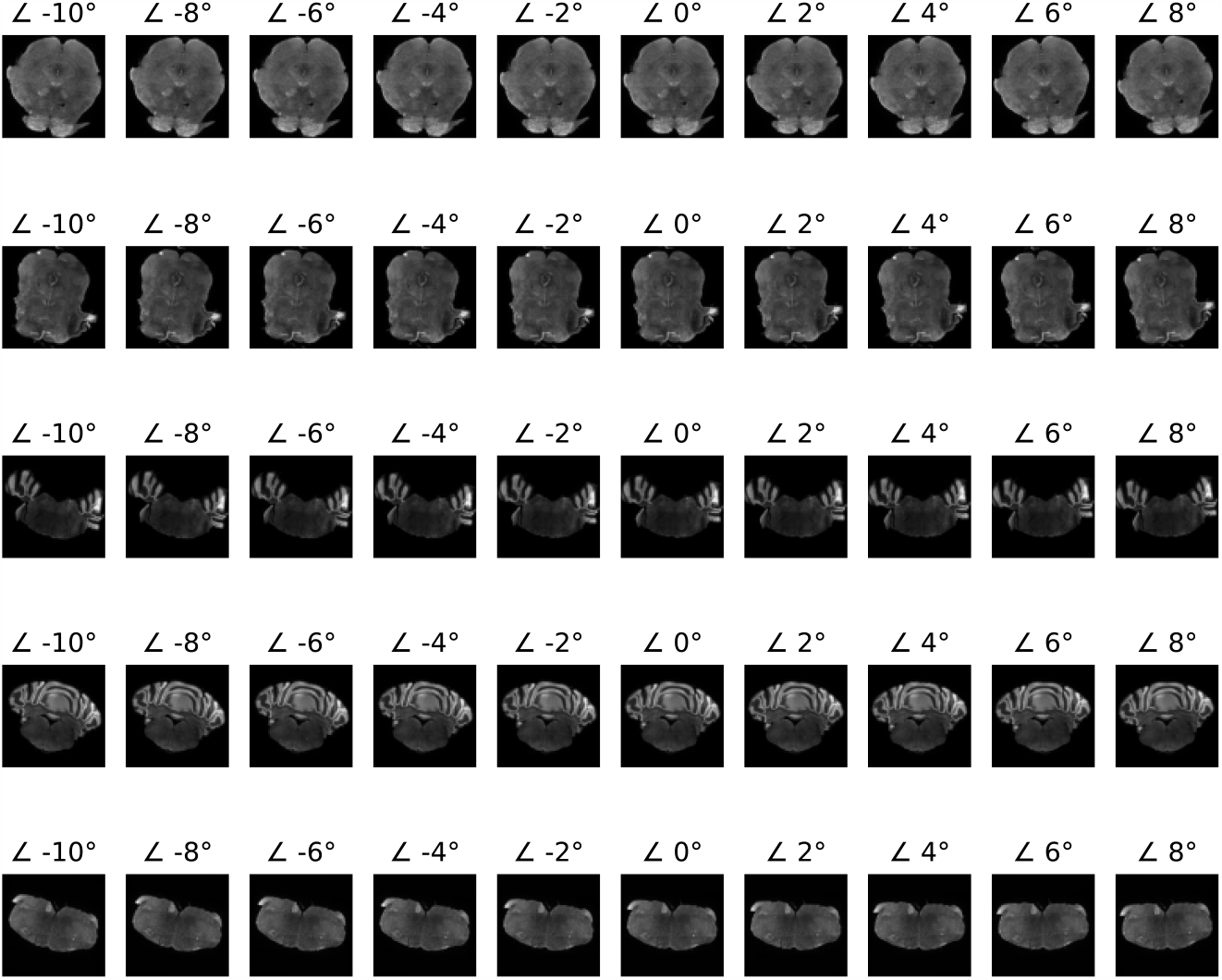
Data augmentation examples are shown on five brainstem sections with rotational variance from -10 to +10 degrees applied on each section.

